# Phylogeography of two Andean frogs: Test of vicariance versus elevational gradient models of diversification

**DOI:** 10.1101/819557

**Authors:** Daria Koscinski, Paul Handford, Pablo L. Tubaro, Peiwen Li, Stephen C. Lougheed

## Abstract

The tropical and subtropical Andes have among the highest levels of biodiversity in the world. Understanding the forces that underlie speciation and diversification in the Andes is a major focus of research. Here we tested two hypotheses of species origins in the Andes: 1. Vicariance mediated by orogenesis or shifting habitat distribution. 2. Parapatric diversification along elevational environmental gradients. We also sought insights on the factors that impacted the phylogeography of co-distributed taxa, and the influences of divergent species ecology on population genetic structure. We used phylogeographic and coalescent analyses of nuclear and mitochondrial DNA sequence data to compare genetic diversity and evolutionary history of two frog species: *Pleurodema borellii* (Family: Leiuperidae, 130 individuals; 20 sites), and *Hypsiboas riojanus* (Family: Hyllidae, 258 individuals; 23 sites) across their shared range in northwestern Argentina. The two showed concordant phylogeographic structuring, and our analyses support the vicariance model over the elevational gradient model. However, *Pleurodema borellii* exhibited markedly deeper temporal divergence (≥4 Ma) than *H. riojanus* (1-2 Ma). The three main mtDNA lineages of *P. borellii* were nearly allopatric and diverged between 4-10 Ma. At similar spatial scales, differentiation was less in the putatively more habitat-specialized *H. riojanus* than in the more generalist *P. borellii*. Similar allopatric distributions of major lineages for both species implies common causes of historical range fragmentation and vicariance. However, different divergence times among clades presumably reflect different demographic histories, permeability of different historical barriers at different times, and/or difference in life history attributes and sensitivities to historical environmental change. Our research enriches our understanding of the phylogeography of the Andes in northwestern Argentina.

## INTRODUCTION

The Tropical Andes hotspot, a region of over 1.5 million km^2^ extending from western Venezuela to northern Argentina, is one of the most biodiverse regions on Earth [1]. This extraordinary diversity includes approximately 30,000 flowering plant species, almost one fifth of known bird species, and some 980 amphibian species of which about 70% are endemic, with habitats spanning alpine grasslands and dry scrub to humid montane forests [2,3]. At its southern terminus this montane forest system exhibits a complex phytogeographic pattern with open microphyllous forests at the base of the Andes, dense Yungas forests at mid-elevations, then alder woodlands grading into alpine meadows, grasslands and steppe (see [4]). Despite the unparalleled species and habitat richness of this region, we know relatively little about the forces that promote or maintain its diversity [5–8], and this deficit is pronounced for the extreme southern portions of the Tropical Andes Hotspot.

Two non mutually-exclusive hypotheses have dominated discussions of speciation and diversification in the Andes: vicariance mediated by orogenesis or shifting habitat distributions, and diversification along elevational environmental gradients [6]. The former hypothesis predicts that sister clades should be distributed allopatrically among adjacent areas that are or were separated by a barrier to gene flow (e.g. [6,9,10]). For the latter, we expect closely related lineages to be parapatrically-arrayed along an elevation gradient (e.g. [11–13]), and for recently arisen gradients we further predict that basal lineages will be at lower elevations than those that are of more recent origin [13,14]. Comparing phylogeographic patterns of co-distributed taxa enhances tests of biogeographical hypotheses [15,16]. For example, spatially and temporally concordant phylogeographic patterns imply that common historical barriers affected past gene flow and produced similar levels and patterns of differentiation [17]. Alternatively, discordant patterns may highlight the importance of differences in ecology and demography [18] or suggest different evolutionary histories [19].

Given their diverse breeding biology and habitats, frogs offer excellent possibilities for examining the factors involved in evolutionary divergence, diversification and speciation. Our focal species, the Rioja treefrog, *Hypsiboas riojanus* [20] (formerly *Hypsiboas andinus* [21]; see [22]) and the rufous four-eyed frog, *Pleurodema borellii* [23], are broadly sympatric in northwestern Argentina and western Bolivia but differ in their life history (see below). Using mtDNA, Koscinski et al. [24] reported well-demarcated clades in *H. riojanus* distributed along a north-south axis (Fig 1), and suggested Pleistocene range fragmentation and subsequent expansion. Here we analyze phylogeographic patterns of *P. borellii* using nuclear and mitochondrial sequences, and add nuclear DNA sequence data and new analytical approaches to our previous analysis of *H. riojanus* mitochondrial sequence data [24]. We compare phylogeographic patterns of *P. borellii* and *H. riojanus* to better understand the roles of vicariance and elevation gradients in clade diversification. We predict spatially and temporally concordant phylogeographic patterns of *P. borellii* and *H. riojanus,* and vicariance as the main driver of clade diversification in both focal species.

**Fig 1.**
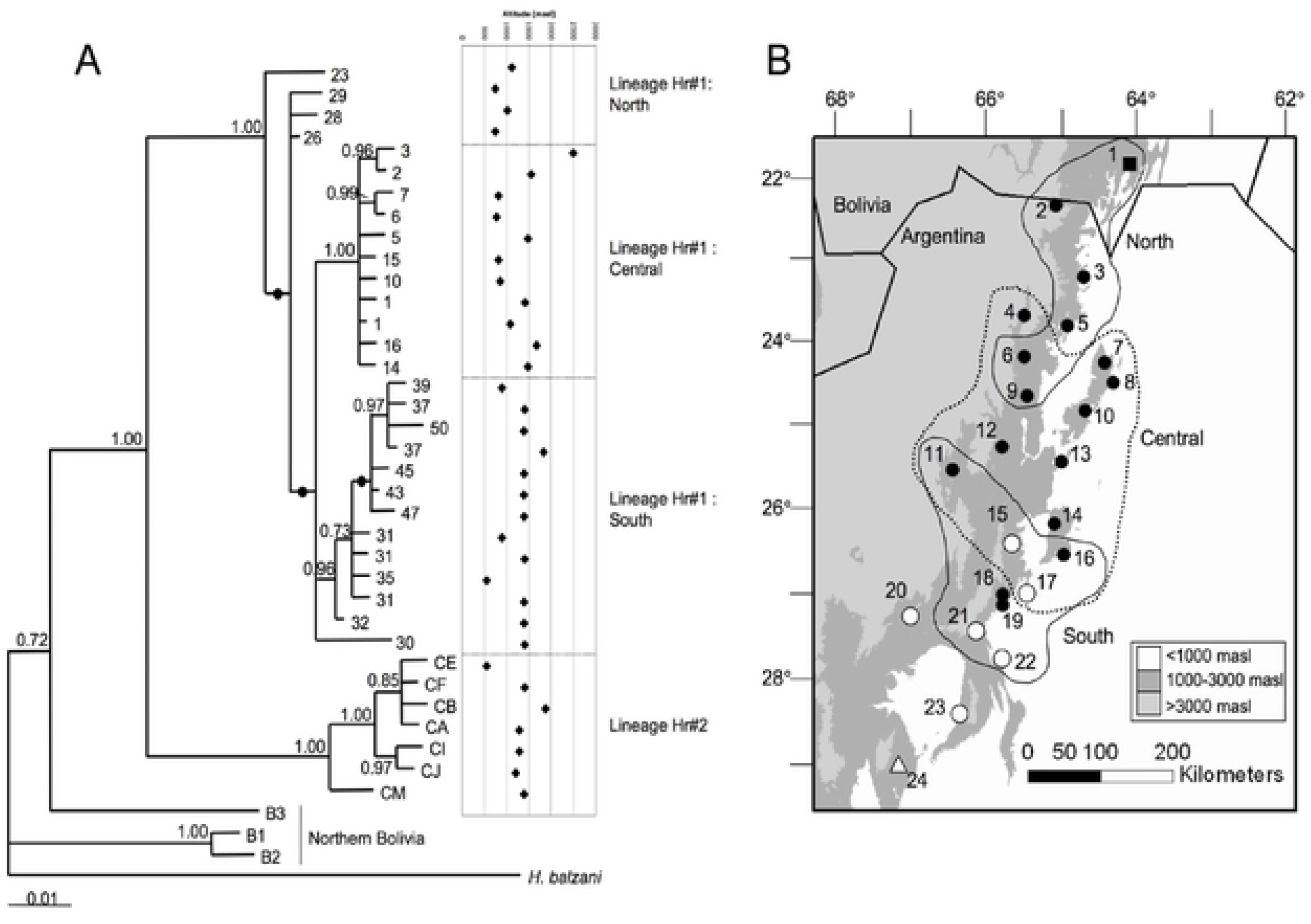
A) Phylogenetic relationships of Argentine and Bolivian *Hypsiboas riojanus* based on Bayesian analyses of cytochrome *b* and control region sequences, using *H. balzani* as an outgroup. Posterior probability values are indicated for each branch, filled circles indicate values less than 0.70. Modified from Koscinski et al. [24]. The elevation of each haplotype is indicated to the right of the tree. B) Geographical distribution of clades from lineage Hr#1 based on control region sequences. Populations within the 3 main groups of lineage Hr#1 that also contain haplotypes from lineage Hr#2 are indicated with unfilled circles (populations #20, 23 and 24 contain only lineage Hr#2 haplotypes). Modified from Koscinski et al. [24]. Locations of sharp genetic discontinuities, likely corresponding to barriers to gene flow among populations are superimposed on the map. Population pair-wise F_ST_ were analysed using Monmonier’s maximum difference algorithm as implemented in Barrier [42]. Analyses were run for one barrier across the whole study area (double line) and then for two barriers (solid lines) across populations within lineage Hr#1 only. Modified from Koscinski et al. [44].

## METHODS

### Study species

*Hypsiboas riojanus* is a moderate-sized (50-60 mm, snout vent length [SVL]) treefrog (family Hylidae) native to the eastern slopes of the Andes Mountains of northwestern Argentina and Bolivia. Previous work showed that northern Bolivian individuals comprise a lineage distinct from those in northwestern Argentina and southern Bolivia [22,24]. We therefore focus solely on Argentina and southern Bolivia. *Hypsiboas riojanus* is mostly found breeding in riparian zones, ditches or flooded areas [25,26, personal observation] spanning lowland montane forests (500 meters above sea level - masl) to montane grasslands (>1500 masl), but is not found in the low elevation (<400 masl) seasonally dry chaco to the east of the mountains nor in high elevation puna (>3400 masl; [26]). *Pleurodema borellii* is a moderate-sized (40-55 mm, SVL) frog (family Leiuperidae) that inhabits open, often disturbed, habitats throughout the moist montane forests and grasslands, but also drier habitats such as the Monte desert, the high elevation puna grass- and scrub-lands, and the western margin of the lowland chaco [25,27]. The high elevation populations in the northernmost part of our sampling area are within the distribution of the sister taxon to *P. borellii*, *P. cinereum*. Molecular studies are inconclusive regarding the distinction of the two putative species [28,29] and thus we include both in our survey.

### Sampling

Samples for this study were either (i) field collected in 1987, 2001, 2004, 2005 or 2006 (*H. riojanus* = 258, *P. borellii* = 126), or (ii) tissues loaned from the Museo Nacional de Ciencias Naturales (MNCN), Madrid, Spain (*H. riojanus* = 6, *P. borellii* = 2), or the Museo de La Plata, La Plata, Argentina (*P. borellii* = 2). Individuals of *H. riojanus* were collected from 23 sites, and of *P. borellii* from 20 sites, across the Argentine portion of the species’ range (Table 1 and Table 2, Fig 1 and Fig 2). The sampling sites for *P. borellii* range from 550 to 3698 masl, and for *H. riojanus* from 550 to 2658 masl. At each site, wetlands (streams, ditches, ponds) were typically searched between the hours of 21:00 and 03:00 local time and individuals located by male advertising calls (females and tadpoles were sampled if encountered). All frogs were captured by hand. Tissues for analysis included toe clips (frogs released), or liver for voucher specimens collected for local museums. All tissue was stored in 70% ethanol until analysis. At site #14 we collected eggs of *P. borellii* to augment sample size. Only one egg per nest was used for analyses. *Hypsiboas balzani* [30] and *P. tucumanum* [31] at one site each were sampled as outgroups. Voucher specimens for *P. borellii* are listed in S1 Table (details for *H. riojanus* in [24]). Voucher specimens were fixed in 10% formalin and then stored in 70% ethanol.

**Table 1.** Sources of tissues and sample sizes for *Pleurodema borellii*. Site numbers refer to Fig 2. Samples were collected by the authors unless otherwise noted.

|  | Site | Elevation<br>(masl) | mtDNA<br>n | Nuclear<br>n |
| --- | --- | --- | --- | --- |
| 1 | Tarija, Bolivia* | 1016 | 2 | 2 |
| 2 | Abra Pampa, Jujuy | 3486 | 5 | 5 |
| 3 | Tres Cruces, Jujuy | 3698 | 1 | 1 |
| 4 | San Andres, Salta | 742 | 8 | 8 |
| 5 | Tropico de Capricornio,<br>Jujuy | 2612 | 5 | 5 |
| 6 | San Francisco, Jujuy | 1588 | 9 | 8 |
| 7 | Tiraxi, Jujuy | 1654 | 15 | 15 |
| 8 | Villa Monte, Jujuy | 1083 | 16 | 16 |
| 9 | Maiz Gordo, Jujuy | 855 | 9 | 9 |
| 10 | Ruta de Cornisa, Salta | 1413 | 9 | 9 |
| 11 | P. N. El Rey, Salta | 913 | 1 | 1 |
| 12 | Pulares, Salta | 1364 | 10 | 9 |
| 13 | Ojo de Agua, Salta | 1108 | 8 | 8 |
| 14 | San Pedro, Tucumán | 940 | 7 | 2 |
| 15 | Yerba Buena, Tucumán† | 513 | 3 | 3 |
| 16 | La Angostura, Tucumán | 1833 | 9 | 8 |
| 17 | Infernillo, Tucumán | 3091 | 3 | 2 |
| 18 | Las Estancias, Catamarca | 1392 | 1 | 1 |
| 19 | Embalse Escaba, Tucumán | 650 | 7 | 7 |
| 20 | San Ignacio, Tucumán | 550 | 2 | 2 |

\* Tissue loaned from the Museo Nacional de Ciencias Naturales, Madrid, Spain.
† Two of the three tissues loaned from the Museo de La Plata, La Plata, Argentina.

**Table 2.**
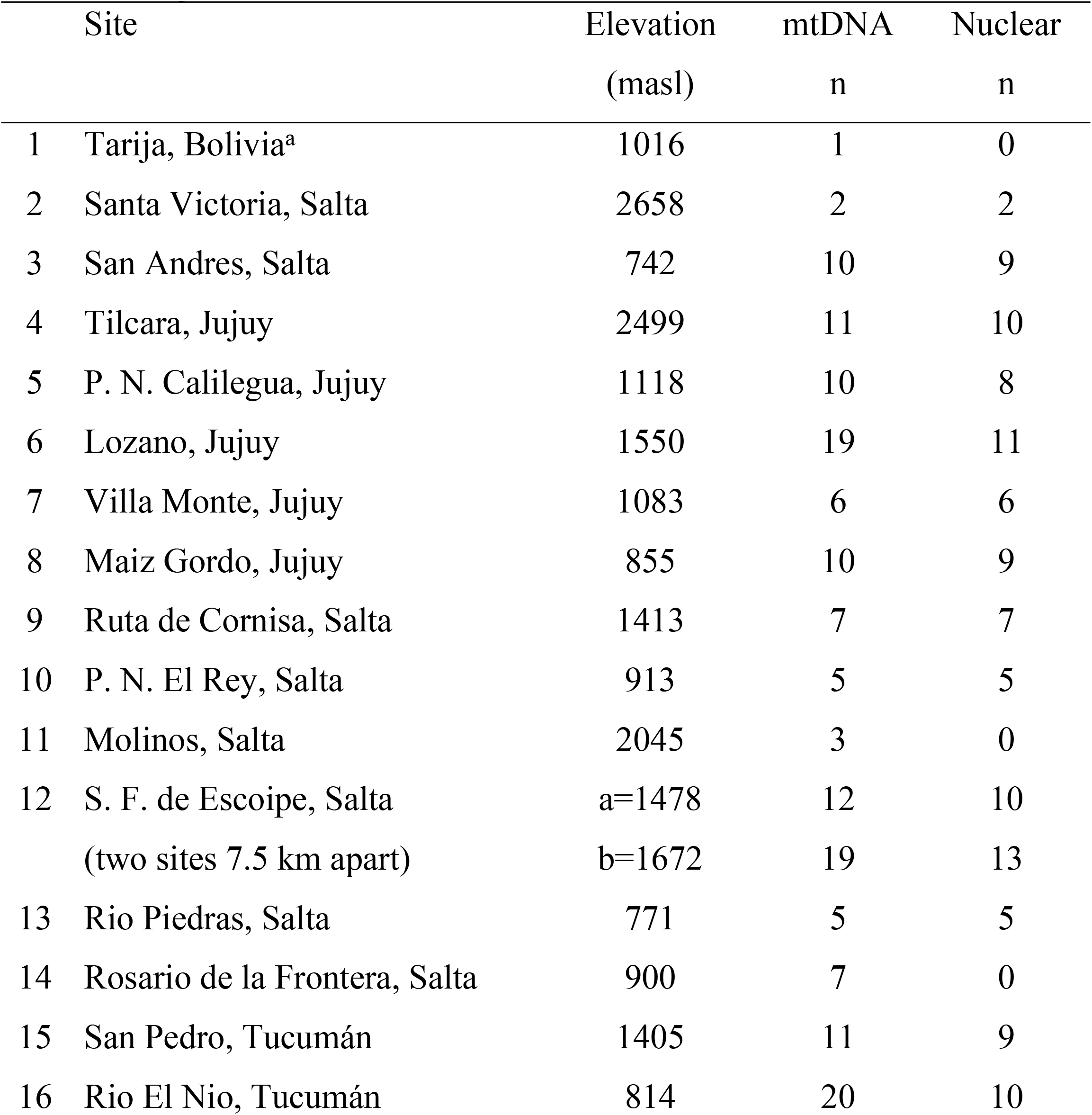
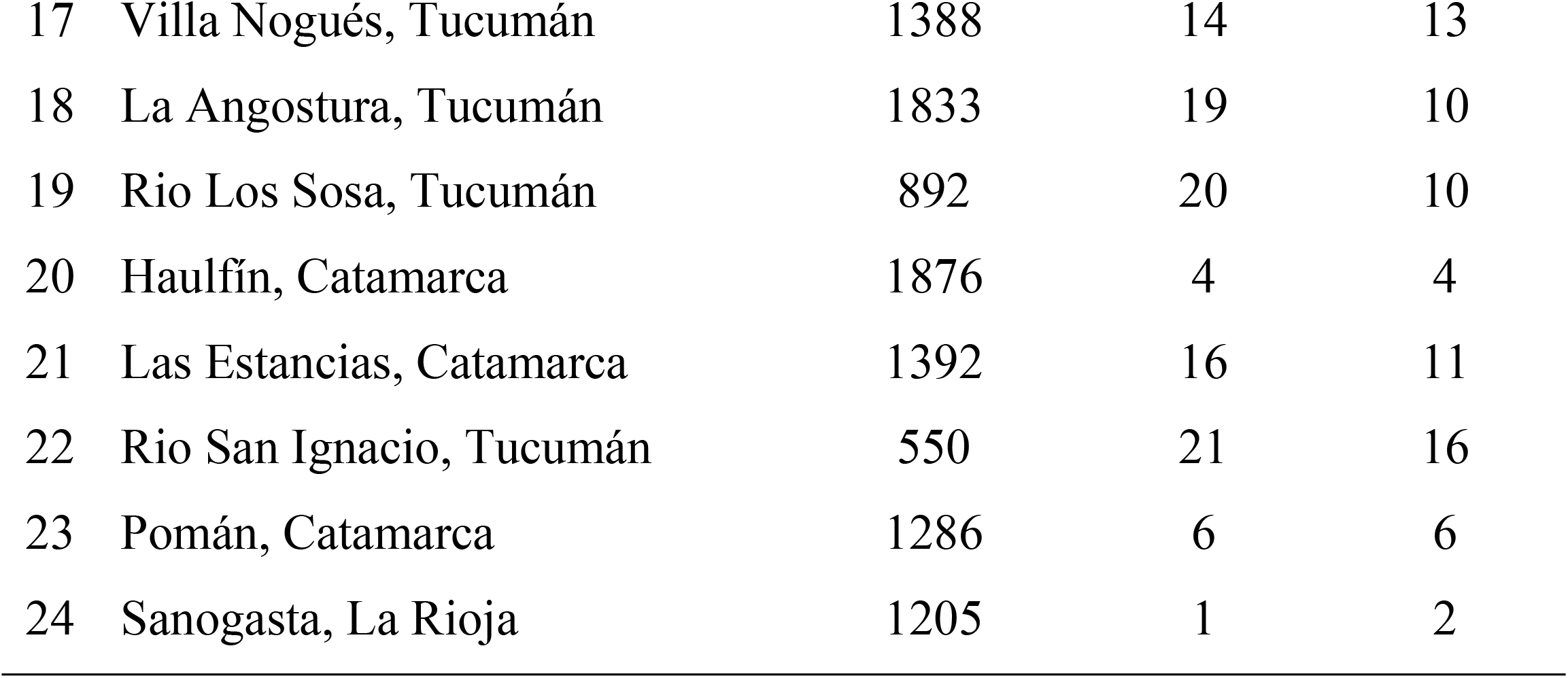
Sample sizes for nuclear analyses of *Hypsiboas riojanus*. Site numbers refer to Fig 1. Sites listed with n = 0 are individuals that were analysed for the mitochondrial fragments [24] but not for the nuclear fragment.

| Site | Elevation<br>(masl) | mtDNA<br>n | Nuclear<br>n |
| --- | --- | --- | --- |
| 1 Tarija, Bolivia<sup>a</sup> | 1016 | 1 | 0 |
| 2 Santa Victoria, Salta | 2658 | 2 | 2 |
| 3 San Andres, Salta | 742 | 10 | 9 |
| 4 Tilcara, Jujuy | 2499 | 11 | 10 |
| 5 P. N. Calilegua, Jujuy | 1118 | 10 | 8 |
| 6 Lozano, Jujuy | 1550 | 19 | 11 |
| 7 Villa Monte, Jujuy | 1083 | 6 | 6 |
| 8 Maiz Gordo, Jujuy | 855 | 10 | 9 |
| 9 Ruta de Cornisa, Salta | 1413 | 7 | 7 |
| 10 P. N. El Rey, Salta | 913 | 5 | 5 |
| 11 Molinos, Salta | 2045 | 3 | 0 |
| 12 S. F. de Escoipe, Salta | a=1478 | 12 | 10 |
| (two sites 7.5 km apart) | b=1672 | 19 | 13 |
| 13 Rio Piedras, Salta | 771 | 5 | 5 |
| 14 Rosario de la Frontera, Salta | 900 | 7 | 0 |
| 15 San Pedro, Tucumán | 1405 | 11 | 9 |
| 16 Rio El Nio, Tucumán | 814 | 20 | 10 |

|  |  |  |  |  |
| --- | --- | --- | --- | --- |
| 17 | Villa Nogués, Tucumán | 1388 | 14 | 13 |
| 18 | La Angostura, Tucumán | 1833 | 19 | 10 |
| 19 | Rio Los Sosa, Tucumán | 892 | 20 | 10 |
| 20 | Haulfín, Catamarca | 1876 | 4 | 4 |
| 21 | Las Estancias, Catamarca | 1392 | 16 | 11 |
| 22 | Rio San Ignacio, Tucumán | 550 | 21 | 16 |
| 23 | Pomán, Catamarca | 1286 | 6 | 6 |
| 24 | Sanogasta, La Rioja | 1205 | 1 | 2 |

**Fig 2.**
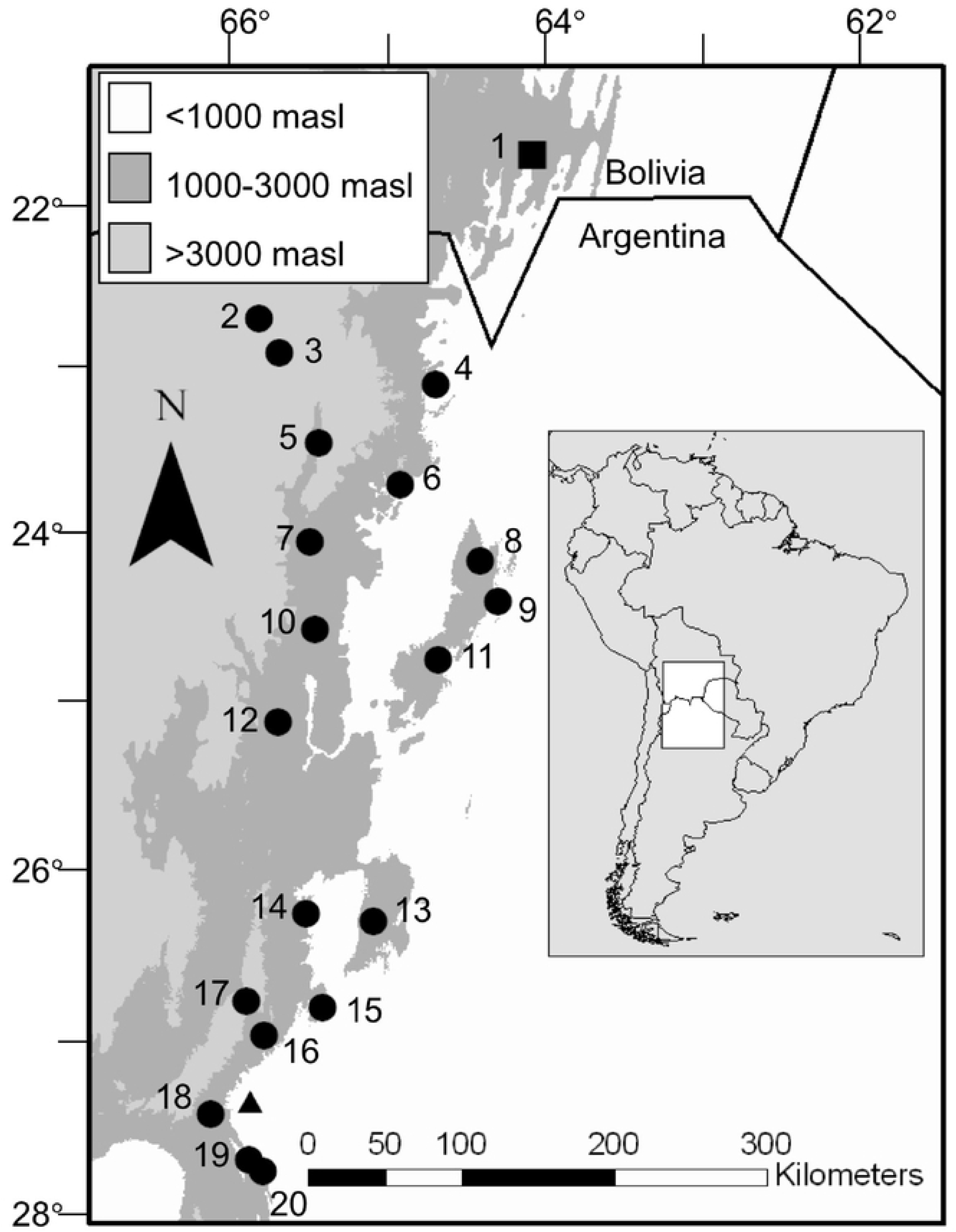
Sampling localities of *Pleurodema borellii* in Northwestern Argentina (circles) and Bolivia (square), and the outgroup taxon *Pleurodema tucumanum* (triangle). Site names and sample sizes for *P. borellii* are listed in Table 1.

### DNA Extraction, PCR and Sequencing

DNA was extracted using standard phenol–chloroform protocols [32] or the DNeasy Tissue Kit (QIAGEN) according to the manufacturer’s protocol. Based on preliminary screening and differences in the number of potentially phylogenetically informative sites among genes, we used different combinations of one nuclear and two mitochondrial markers for each focal species. For *H. riojanus* we combined portions of the mitochondrial control region (663 base pairs [bp]), 358 bp of cytochrome *b* (cyt *b*), and 380 bp of the exon of the nuclear recombination activating 1 (RAG1) gene, and for *P. borellii* we used 538 bp of mitochondrial 16S rDNA sequence, 380 nt of cyt *b*, and 316 bp fragment of the nuclear rhodopsin exon 1 sequence (RHOD). PCR and sequencing methods are provided in S1 Text. Homologous sequences were aligned in Clustal X, ver. 1.83 [33].

Allelic variants of heterozygotes from nuclear genes were inferred using fastPHASE version 1.2 [34] with default settings. Cloning and sequencing of PCR fragments were used to confirm alleles in several individuals of *H. riojanus* (see S1 Text). We used Genepop web version 4.0.10 [35,36] to estimate allele frequencies within each sampled population (defined as comprising samples from within a 10 km radius) and to test for departure from Hardy-Weinberg expectations using the Markov Chain method with default settings.

### Phylogenetic and Phylogeographic Analyses

We combined phylogenetic and network approaches to infer phylogeographic patterns because initial surveys revealed substantial intraspecific phylogenetic diversity. For *H. riojanus*, we previously generated both Bayesian and maximum likelihood (ML) phylogenetic trees using combined control region and cyt *b* sequences, with *H. balzani* as an outgroup [24]. To allow comparison of mitochondrial genealogical histories, for *P. borellii* we also conducted phylogenetic analysis of a combined data set of cyt *b* and 16S for all samples with *P. tucumanum* as an outgroup.

We previously constructed a haplotype network for *H. riojanus* control region [24] and here add one for RAG1. For *Pleurodema borellii*, we also constructed separate networks for the combined mtDNA data set (cyt *b* and 16S) and the nuclear RHOD sequence data. Statistical Parsimony Networks were generated in TCS version 1.21 [37]. Recognizing recent criticisms of the mismatch distribution analysis [38], but also the many studies that deploy it, we examined recent demographic patterns using two approaches. We performed mismatch distribution analyses for each main clade within species, and tested against a model of sudden expansion using the generalized non-linear least-squares approach [39]. The validity of the model was evaluated using 1000 parametric bootstraps in Arlequin v. 2.000 [40]. We also used a coalescent approach and Bayesian skyline plots [41] of cyt *b* data to infer demographic history of each species. To investigate locations of potential barriers to gene flow we used Monmonier’s maximum difference algorithm as implemented in Barrier v. 2.2 [42,43]. We compared the results for *H. riojanus* [44] to those for *P. borellii* to see if phylogeographic break points were spatially coincident. We estimated time to most recent common ancestor (TMRCA) for major clades for each surveyed species using cyt *b* sequences using BEAST v. 1.4.6 [45]. Greater detail on phylogenetic analyses is provided in S1 Text.

## RESULTS

A total of 535 bp sequence of 16S and 364 bp of cyt *b* was obtained from 130 individuals of *P. borellii*. The *H. riojanus* dataset comprised the aforementioned 340 bp of the control region from 258 individuals plus a 358 bp fragment of cyt *b* from a subset of 40 individuals representing unique control region haplotypes (see [24]). All mitochondrial sequences showed an overall AT nucleotide bias that is characteristic of vertebrate mtDNA [46], plus the biases towards transitions and third positions in cyt *b* expected of a protein-coding gene. Neither stop codons nor indels were found in the cyt *b* fragments.

We obtained 285 bp of RHOD from 120 individuals of *P. borellii*, and 303 bp of RAG1 from 185 individuals of *H. riojanus*. No stop codons were found in these sequences. Seven variable sites defined 10 RHOD haplotypes and 10 variable sites defined 13 RAG1 haplotypes. One population of *P. borellii* (site #16) exhibited heterozygote deficiency after sequential Bonferonni correction [47]. No significant departures from Hardy-Weinberg equilibrium were detected in *H. riojanus*. The individuals used as outgroups of both species were heterozygotes containing haplotypes not found in the ingroup.

### Phylogeographic patterns for *P. borellii*

The model of evolution selected for *P. borellii*, using AIC for the combined data set of cyt *b* and 16S, was the general time-reversible model with a proportion of invariable sites, and gamma-distributed rates (GTR + I + G). Trees using ML and Bayesian methods recovered similar topologies, with generally stronger support from Bayesian analyses.

Three well-supported mtDNA lineages (Fig 3) are distributed north to south with little geographic overlap and equivocal evidence for parapatric distributions along elevational gradients in one lineage. These lineages corresponded to three separate statistical parsimony networks that could not be connected at the 95% parsimony level (Fig 4). For consistency with the phylogenetic results we refer to each network as a lineage. Lineage Pb#1 and lineage Pb#2 were separated by at least 23 steps, and lineage Pb#1 and Pb#3 by 27 steps. The most divergent lineage (Pb#3) contained solely individuals collected in northern Jujuy and southern Bolivia (sites #1, 2, 3 and 5, Fig 5), and may represent *P. borellii*’s putative sister taxon, *P. cinereum*. The elevational distribution within lineage Pb#3 suggests an elevational effect, as its southern Bolivian haplotypes are found at much lower elevations than all other haplotypes. Lineages Pb#1 and Pb#2 showed an unusual geographic distribution. Lineage Pb#2 was found only in the central portion of the range, whereas lineage Pb#1 was found in the northeast and southwest, but not centrally. All three clades within lineage Pb#1 were mutually allopatric. Lineages Pb#2, Pb#3, and all three main clades in lineage Pb#1 exhibited patterns characteristic of rapid population expansion using mismatch distributions.

**Fig 3.**
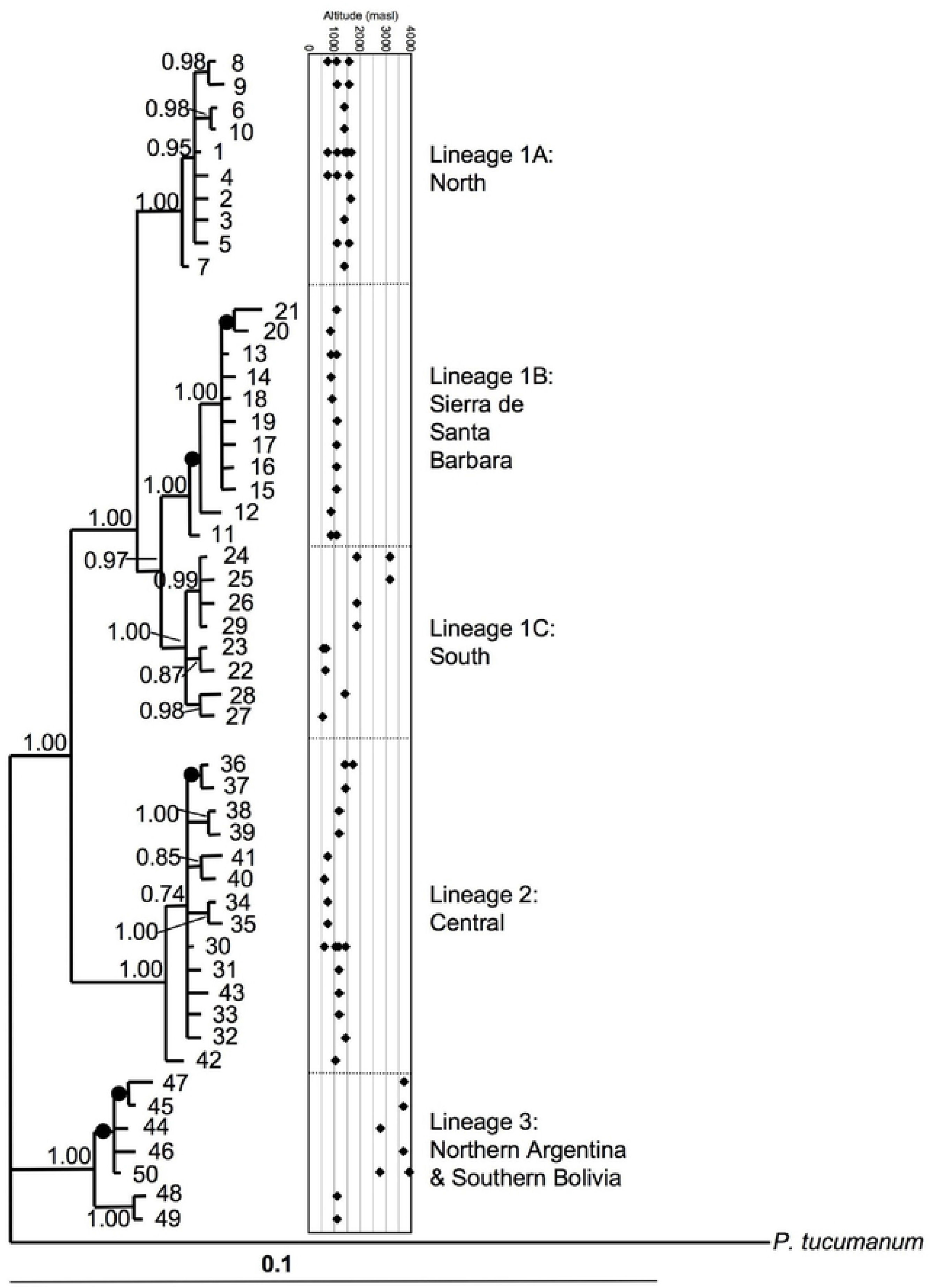
Phylogenetic relationships of Argentine and Bolivian *Pleurodema borellii* based on Bayesian analyses of 16S and cytochrome *b* DNA sequences, using *P. tucumanum* as an outgroup. Posterior probability values are indicated for each branch, filled circles indicate values less than 0.70. Haplotype codes correspond to those shown in Fig 4. The elevation of each haplotype is indicated to the right of the tree, and haplotypes can be found at multiple localities.

**Fig 4.**
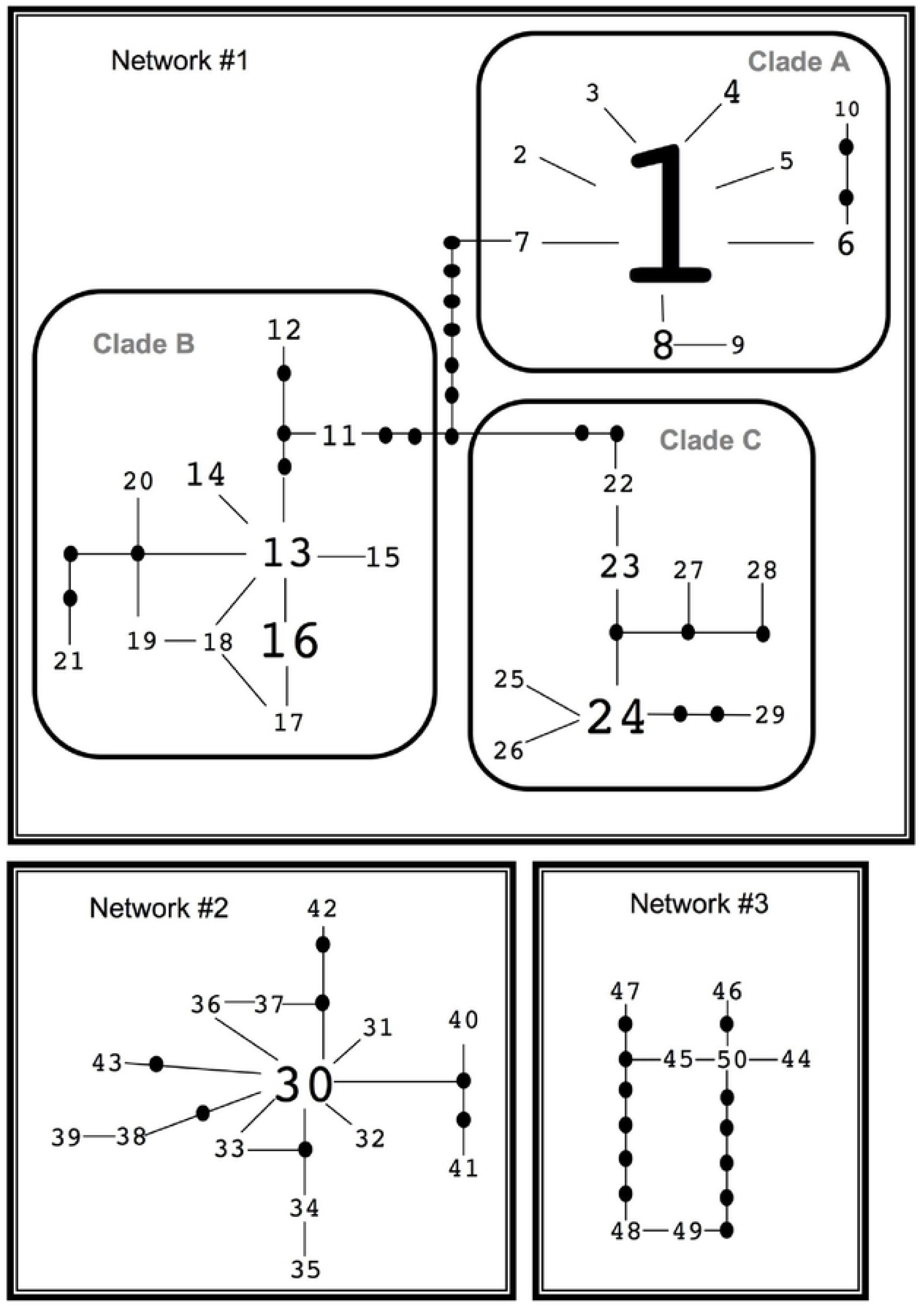
Maximum parsimony networks using mtDNA 16S and cytochrome *b* genes of *Pleurodema borellii*. Filled circles indicate missing or unsampled haplotypes and font size approximates relative sample size for each haplotype. The three networks could not be connected at the 95% parsimony level.

**Fig 5.**
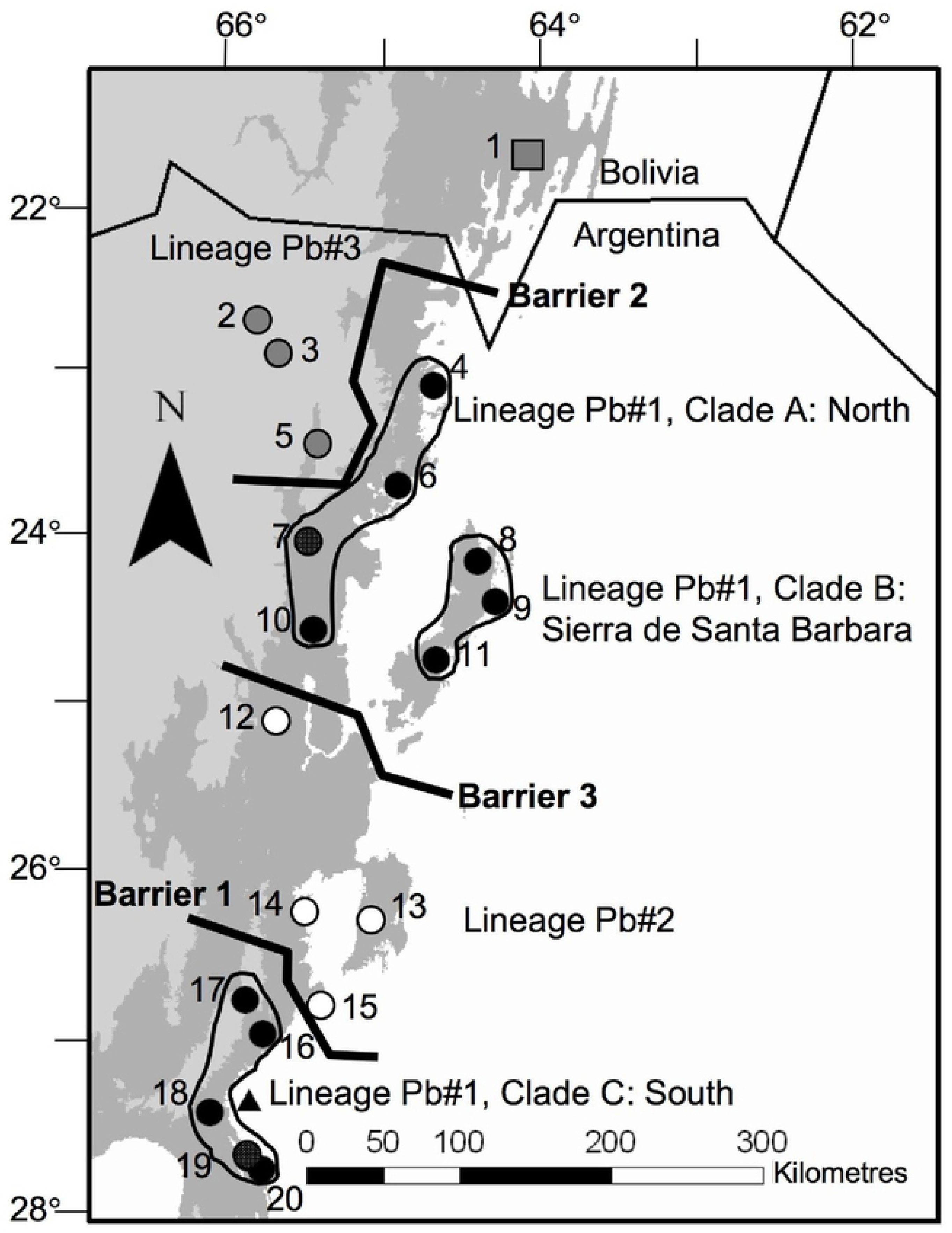
Geographical distribution of mtDNA lineages within *Pleurodema borellii*. Populations indicated with the same color belong to the same lineage: black = lineage Pb#1 with main clades grouped using solid black lines, white = lineage Pb#2, grey = lineage Pb#3. Populations #7 and #19 are indicated with black and white circles as each population contains individuals belonging to lineage Pb#1 and Pb#2. Locations of sharp genetic discontinuities (Barriers 1, 2 and 3), likely corresponding to barriers to gene flow among populations of *Pleurodema borellii*. Population pair-wise F_ST_ were analysed using Monmonier’s maximum difference algorithm as implemented in Barrier [42]. Shading represents elevation as in Fig 2.

We identified three barriers using Monmonier’s maximum difference algorithm with mtDNA data for *P. borellii* that correspond well with the three main identified lineages identified from our phylogenetic analysis (Fig 5)

The RHOD network showed two clusters of haplotypes connected by a single unsampled or extinct haplotype (Fig 6). The two most common haplotypes in *P. borellii* differed by four substitutions, and were distributed throughout the study area. The remaining haplotypes were found in one to four sites (Fig 6). Nine sites across the study area were fixed for the most common haplotype (rhod-2) and a tenth was nearly fixed (frequency of 0.94) (Fig 6). Site #12 in west-central Salta province contained the highest number of haplotypes (7 of 10 ingroup haplotypes) even though other populations had equal or greater sample sizes. The same locality was most diverse for *H. riojanus* RAG1 alleles (see below). The geographic distribution of RHOD haplotypes does not show the striking delineation of mtDNA clades.

**Fig 6.**
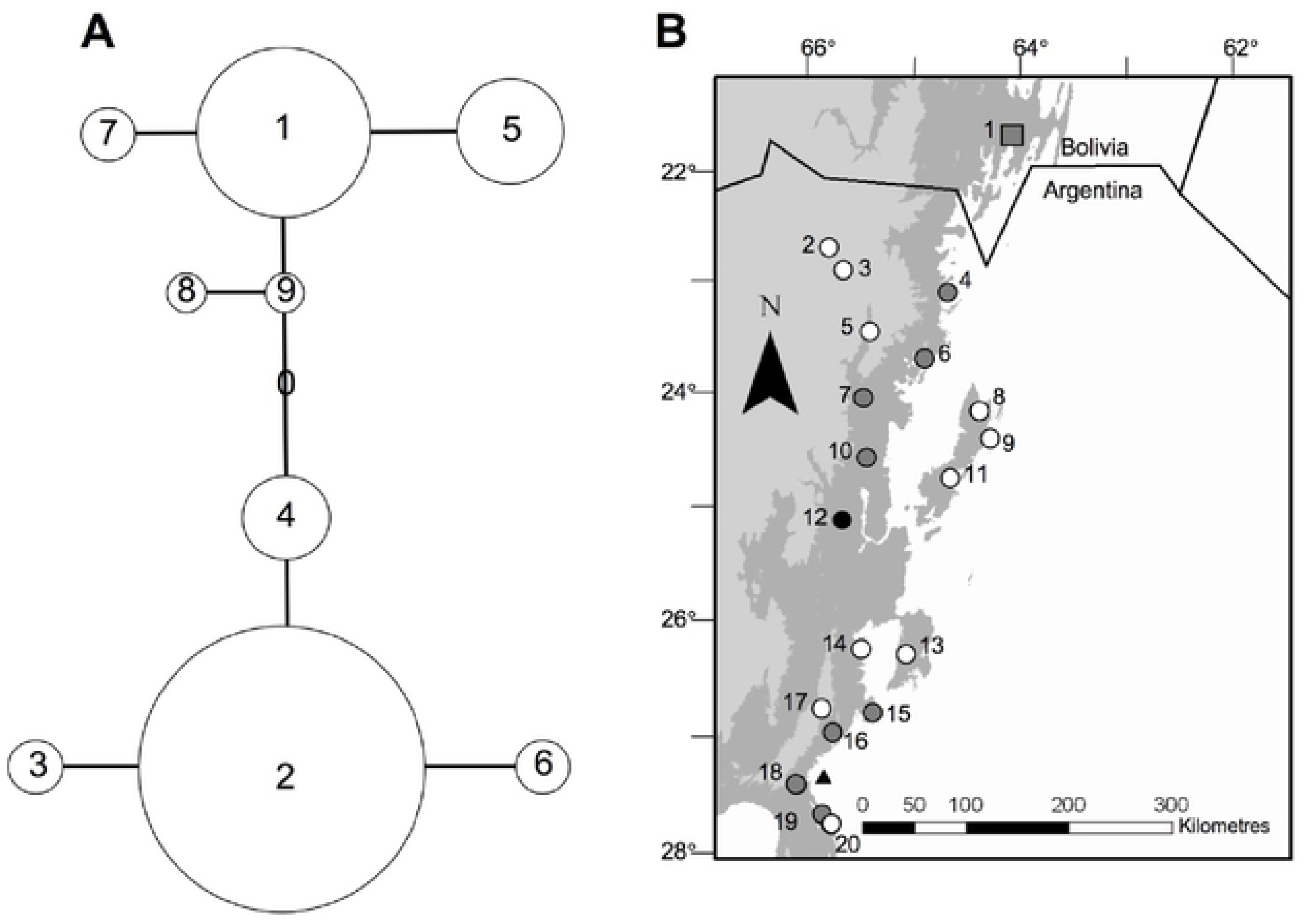
A) Maximum parsimony networks using the rhodopsin gene of *Pleurodema borellii*. Open circle indicates missing or unsampled haplotype. Circle size approximates relative sample size for each haplotype. B) Geographic distribution of population allele frequencies based on rhodopsin genotypes. White = populations fixed for allele rhod-2 (population #9 frequency of 0.944). Gray = populations containing mixtures of haplotypes. Black = population #12 which contains 8 of the 9 haplotypes.

### Phylogeographic patterns for *H. riojanus*

Koscinski et al. [24] found three major mtDNA lineages in *H. riojanus*: one representing northwestern Argentina and southern Bolivia; one from the southern portion of the range; and one representing northern Bolivia (Fig 1). Recent evidence suggests that *H. riojanus* from northwestern Argentina and southern Bolivia – formerly referred to as *H. andinus* – and *H. riojanus* are synonymous [22] and we treat them as a single species (*H. riojanus*) here. The divergent northern Bolivian populations were excluded from this study. Within the geographically extensive *H. riojanus* lineage (Hr#1), we found three clades distributed along a north-south axis (Fig 1).

Analyses of *H. riojanus* RAG1 data generally support findings from mitochondrial DNA. The network for RAG1 contained ambiguous connections and resembles a starburst, with many haplotypes closely related to a central, common haplotype (Fig 7). The two most common RAG1 haplotypes were distributed widely but many other haplotypes were unique to one or two populations. Most populations contained two to four haplotypes. A single site (#12) contained 8 of the 13 haplotypes. This locality is within 30 km of the site that was most diverse for *P. borellii* RHOD alleles. Haplotype RAG1-5 is only present in northern populations (Fig 7), and is spatially coincident with the Northern mtDNA clade and the northern populations of the Central mtDNA clade (within lineage Hr#1). The southern populations contain only the two common haplotypes (RAG1-1 and RAG1-3).

**Fig 7.**
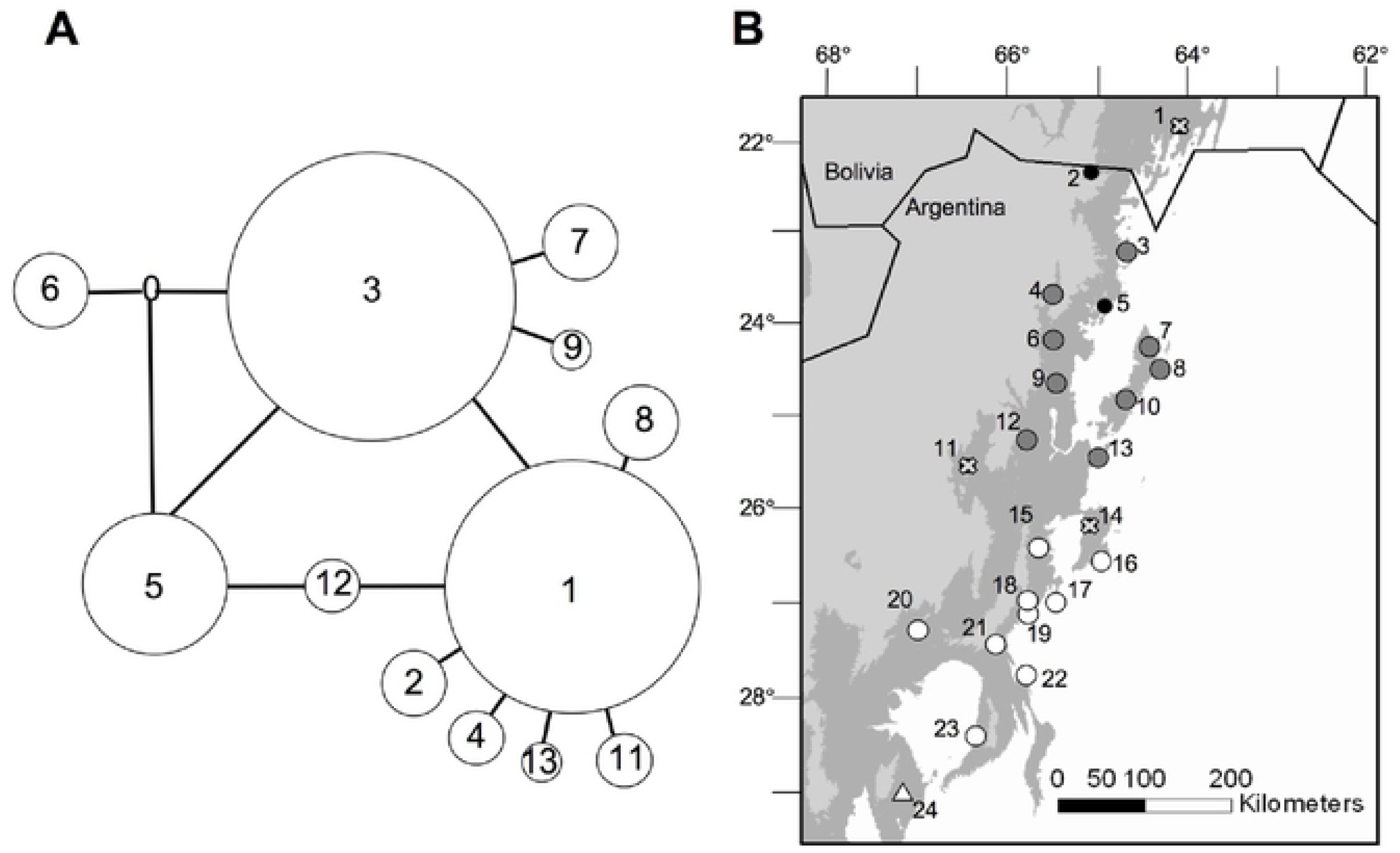
A) Maximum parsimony networks using the RAG1 gene of *Hypsiboas riojanus*. Open circle indicates missing or unsampled haplotype. Circle size approximates relative sample size for each haplotype. B) Geographic distribution of RAG1 haplotypes. Gray = populations containing haplotype RAG1-5, white = populations only containing haplotypes RAG1-1 and RAG1-3, black = mixture of haplotypes but excluding RAG1-5. Localities indicated with an “x” were not included in the analyses.

### Demography & Timing of Divergence

Cyt *b* skyline plots were similar in both species, and the 95% highest posterior density (HPD) intervals overlapped substantially (S1 Fig). Recent (<1 Ma) population expansion is evident in both species, supporting findings using mismatch distributions (S2 Table). To make direct comparisons of the two taxa, we estimated TMRCA for major clades using cyt *b*. In *H. riojanus*, the deepest split (between lineage Hr#1 and lineage Hr#2, was estimated at 2-6 Ma [24]). Within lineage Hr#1, TMRCAs were all less than 2 Ma, placing them within the Pleistocene. The deepest divergences among the main lineages of *P. borellii* (Pb#1, Pb#2 and Pb#3) ranged from 4.2-10 Ma (Table 3). Within lineage Pb#1 of *P. borellii*, the three clades appeared to have diverged between 2-4 Ma. Thus, most divergence events in *P. borelli* in Argentina seem to have occurred in the late Pliocene (>2 Ma), predating those within *H. riojanus*.

**Table 3.**
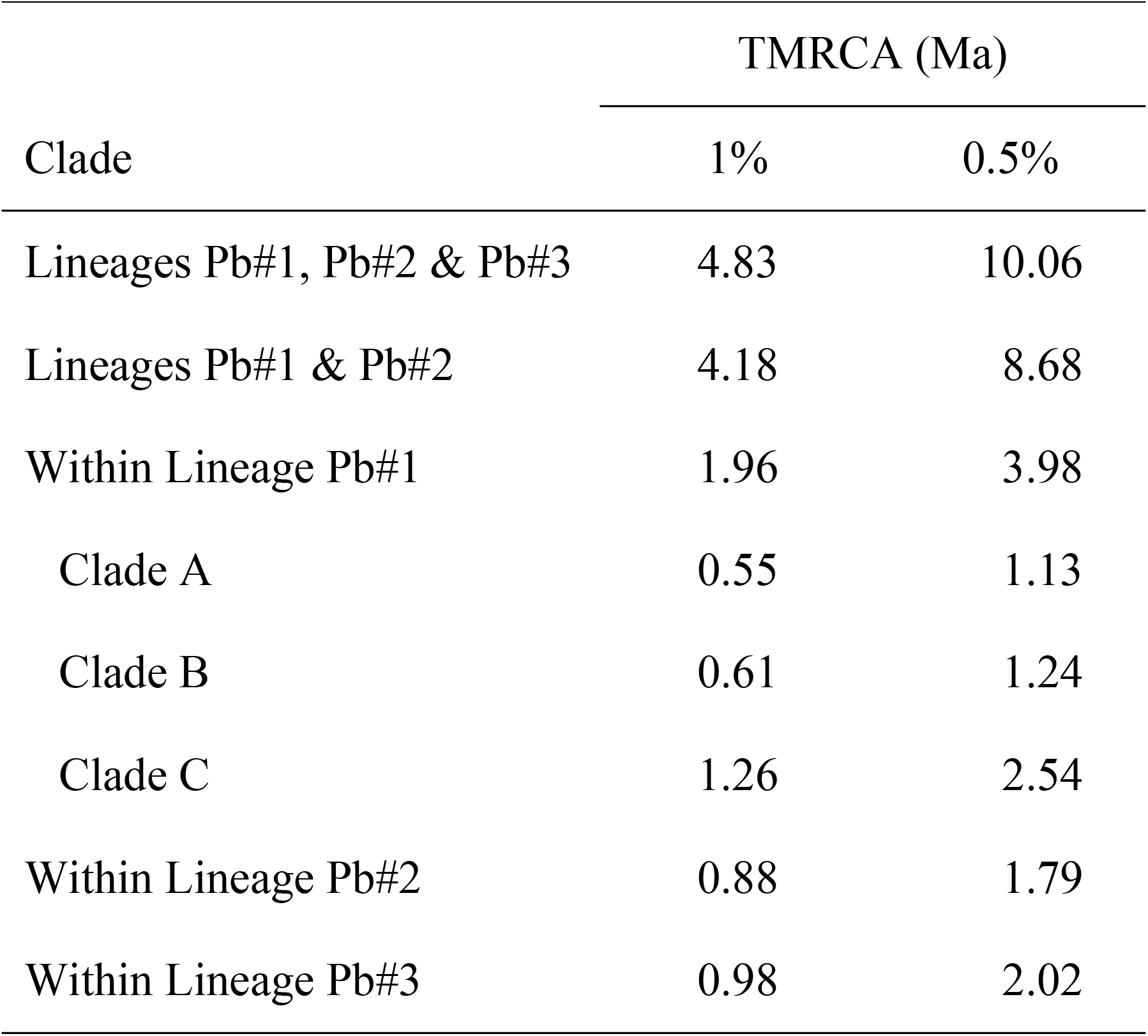
Estimates of divergence time for mitochondrial lineages of *Pleurodema borellii* based on analyses using BEAST (see text for details). The analyses were run using a molecular clock of 1% or 0.5% per million years for cytochrome *b*. Refer to Fig 5 for clade numbers. TMRCA = time to most recent common ancestor, Ma = million years ago.

| Clade | TMRCA (Ma) |  |
| --- | --- | --- |
|  | 1% | 0.5% |
| Lineages Pb#1, Pb#2 & Pb#3 | 4.83 | 10.06 |
| Lineages Pb#1 & Pb#2 | 4.18 | 8.68 |
| Within Lineage Pb#1 | 1.96 | 3.98 |
| Clade A | 0.55 | 1.13 |
| Clade B | 0.61 | 1.24 |
| Clade C | 1.26 | 2.54 |
| Within Lineage Pb#2 | 0.88 | 1.79 |
| Within Lineage Pb#3 | 0.98 | 2.02 |

## DISCUSSION

Shifts in connectivity among populations play important roles in shaping diversification among evolutionary lineages. *Hypsiboas riojanus* and *Pleurodema borellii* show striking phylogeographic structuring across their respective Argentine ranges. For both species, the spatial distribution of mitochondrial clades is consistent with a vicariance model of diversification rather than the environmental gradient model. However, distinct TMRCA estimates for the two species may reflect different demographic/colonization histories, the action of long-standing historical barriers at different times over the late Pliocene and Pleistocene, or difference in life history attributes and sensitivities to historical environmental change. We explore these issues in detail below.

### Lineage Divergence and Phylogeography within *P. borellii*

The north-south disjunct distributions of lineages within *P. borellii* and lack of correspondence of phylogeny with elevation strongly support the vicariance model over the gradient model of divergence. We found three mitochondrial lineages in *P. borellii* in northwestern Argentina and southern Bolivia distributed on a north-south axis. These lineages, however, were not mirrored by patterns in RHOD although all populations were clearly more closely related to each other than to the outgroup *P. tucumanum* for both classes of loci. Evolution of nuclear protein coding genes is slower than that of mtDNA due to larger effective population sizes and thus longer times to reciprocal monophyly [48], so mtDNA divergence would relate to demographic events in the more recent history of the group. Many populations across the range and across divergent mtDNA clades were fixed for haplotype Rhod-2, possibly reflecting connectivity in the deeper past, ancient founder events, or a signature of selection.

The three deeply-diverged *P. borellii* mitochondrial lineages are nearly allopatric suggesting a long history of isolation. We estimate that these lineages separated about 4-10 Ma, a time of rapid uplift of the Andes and formation of novel arid high elevation habitats [49]. This orogeny substantially altered atmospheric circulation and precipitation patterns [49,50], as well as hydrology [50]. Concomitant landscape changes may have significantly affected population connectivity perhaps leading to the divergence of the lineages that we found. Lineages Pb#2 and Pb#3 show mismatch distribution patterns consistent with population growth. If such growth occurred during range expansion, then secondary contact of these divergent lineages would ultimately occur. Consistent with such a supposition, lineage Pb#2 is currently sympatric with lineage Pb#1 at the northern and southern edges of its range across our study area (sites #7 and #19). Mitochondrial lineage Pb#3 occurs in the northernmost part of our study area corresponding with the distribution of the putative sister taxon *P. cinereum* (see [25,28]; [51] for discussion). These lineages may indeed represent cryptic taxa, although all divergences within our samples are much lower than to the outgroup taxon *P. tucumanum*. The differentiation of *P. cinereum* has long been suggested to be due to adaptation to high elevations [25]. Our mtDNA phylogeny groups individuals from high elevation northwestern Argentina populations with low elevation southern Bolivian individuals. Differentiation within this lineage may have been driven by elevational gradients but we can offer no definitive statement here. Further analyses with additional Bolivian samples are needed to evaluate phylogenetic, morphological and acoustic patterns.

Mitochondrial lineage Pb#1 contains three entirely allopatric clades. Given that sister clades B and C are separated by more than 200 km (by territory where only individuals belonging to lineage Pb#2 are found), this pattern is likely due to past fragmentation as long-distance dispersal is unlikely. The three clades originated between 1-7 Ma implying that the events that caused their divergence predate Pleistocene climatic cycling potentially involving Andean uplift. For example, clade B maps to the isolated Sierra de Santa Barbara suggesting historical isolation within this mountain range (Fig 5). This further supports vicariance as the primary driver of differentiation in the group.

Two locales (#7 and 19) showed admixture of mitochondrial lineages Pb#1 and Pb#2. Mismatch distributions suggest population growth in lineage Pb#2 as well as the north and south clades of lineage Pb#1, observations that are consistent with range expansion leading to secondary contact. The geographic distribution of these lineages is puzzling because genetic exchange appears to have continued among the clades of lineage Pb#1 but not between it and the central lineage Pb#2, despite its central location. This “leap-frog” pattern may indicate that unsampled populations link the clades of lineage Pb#1, or that the range of lineage Pb#1 was once more extensive but was displaced by lineage Pb#2. The genetic divergence between these two lineages is nearly 5% at cyt *b*, implying long-term isolation.

### Phylogeography of Northwestern Argentina

Both frog species show strong genetic structure and similar phylogeographic patterns across the study area. Coalescent analyses assuming the same substitution rate suggest that TMRCA for *H. riojanus* lineage Hr#1 is 1-2 Ma, and that for *P. borellii* lineage Pb#1 is 2-4 Ma. Alternatively, we must postulate a two-fold molecular rate difference between the species to make the phytogeographic patterns of both species temporary concordant. Body size, generation time, population size, metabolic rate, and clutch size all can affect the rate of molecular evolution [52–54]. *Hypsiboas riojanus* eggs are deposited in masses of approximately 600 eggs [Marcos Vaira, unpublished] while *Pleurodema borellii* typically deposits between 1000 and 2500 eggs [Marcos Vaira, unpublished; 55: mean for 11 clutches = 1277+/−352]. Generation time for *H. riojanus* is likely one year, similar to the closely related *H. pulchellus* [56]. Generation time has not been estimated for *P. borellii*. Breeding in *P. borellii* is concentrated in the wettest part of the year [55,57] and likely results in a generation time of at least one year, as juveniles that emerge must wait until the next summer to breed. Although the two species differ in fecundity, they are similar in body size, generation time, and likely metabolic rates hence a two-fold difference in mutation rates is unlikely [52,54].

Similar geographic patterns with different temporal signatures may be due to differences in demographic history of each species. Large or highly structured populations tend to show deeper lineage separation in response to a barrier because (i) more ancestral polymorphisms were available which might be fixed in each of the sundered lineages, or (ii) already differentiated and spatially structured lineages may characterize the sundered lineages [58]. Effective population size appeared to be comparable for both species but genetic structure may have differed. Local extinction and recolonization events across the barrier, especially if the barrier is intermittent (e.g. due to climatic cycling) may also produce such patterns [58]. If the barrier formed slowly, or the species’ responses varied, the barrier may have been more effective (and acted longer) for one species than the other. Such events could erase the phylogenetic signal of previous differentiation events, leading to apparent differences in temporal patterns.

*Hypsiboas riojanus* has lower fecundity and a reported limited ability to move away from the riparian zones of its breeding streams [25,26, personal observation]. *Pleurodema borellii* is evidently an opportunistic breeder in diverse habitats, found in high densities when rains fill shallow depressions. Regardless, population connectivity appears to be higher in *H. riojanus*, even across habitats that one would suppose are barriers for a treefrog that relies on water, such as the semi-arid scrubland separating the Sierra de Santa Barbara (*P. borellii* sites #7, 8, 11, Fig 5) from all other populations. This result is counter-intuitive given that several generalist species have been found to display higher gene flow than habitat-restricted species [59–61]. Factors other than fecundity and habitat specialization must therefore play a critical role in driving differentiation and deserve further investigation (e.g. philopatry, sex-biased dispersal).

The differences in timing of these deepest divergence events in the two taxa may be attributable to their different biogeographic histories. The 2-6 Ma date for *H. riojanus* may relate to the recent dispersal of the entire Andean species group into the Andes from the Atlantic Forest of eastern Brazil where the taxon originated [62,63]. Connections between the Atlantic Forest of southeastern Brazil and the Amazon lowland forests have been suggested based on floristic analyses [64] and been shown for terrestrial species [65: small mammals; 66: plants; birds 67–69], as well as obligate freshwater species [70: characiform fish] in the last 10 Ma. The major dispersal route was likely through the Paraná River basin, which drains large areas of the Andes of northwestern Argentina, Paraguay and southern Brazil. There are no molecular phylogenies for *Pleurodema* and analyses with limited samples (mtDNA 16S sequences for six of 12 species, data not shown) do not support the relationships based on morphological characters [71], which suggested a southern Andes origination for the group. If Duellman and Veloso [71] are correct, the deep genetic divergence reported here for *P. borellii* lineages imply that the genus has likely existed in northwestern Argentina for much longer than 10 Ma.

Genetic discontinuities in both species suggest some common areas of genetic divergence, and possible locations of barriers to gene flow in this region. The best-supported gene flow barrier (using mtDNA) lies near ~27°S (see Fig 1 for *H. riojanus*, Fig 7 for *P. borellii*), although no obvious geological, hydrological or habitat disjunctions exist there currently. The divergence among populations in this area dates to approximately 2-6 Ma in *H. riojanus* (lineages Hr#1 and Hr#2) and 5-10 Ma in *P. borellii* (clade 3-3 of lineage Pb#1 and lineage Pb#2).

Recent work on humid montane avifauna has identified the southern Andean Yungas of northwestern Argentina and southern Bolivia as a distinct biogeographic unit, with ties to the Brazilian Atlantic and northeastern Argentine forest systems [72]. Phylogenetic studies of cacti also support the uniqueness of this region for the arid habitats of the Andes [73,74]. We know of four detailed molecular studies that are available for comparison. Clemente-Carvalho et al. [75] examined phylogeographic patterns of the Yungas Redbelly Toad, *Melanophryniscus rubriventris*, in Northwestern Argentina. They found two well-supported intraspecific lineages, one confined to extreme northern Argentina and the other extending southward into Salta province, a pattern that they interpreted as resulting from historical range fragmentation and subsequent expansion driven by Milankovitch cycles. Braun et al. [76] investigated the phylogenetic relationships of sigmodontine rodents (*Akodon* sp.) of Paraguay, Bolivia and Argentina using cytochrome *b*. They showed a north to south distribution of three species from southern Bolivia through northwestern Argentina whose boundaries match the distribution of clades found in our study. Although the phylogeographic history of these rodents is not known, Braun et al. [76] implicate Pliocene and Pleistocene vicariant events as important for origination of the species. Quiroga & Premoli [77] studied a cold-tolerant tree (*Podocarpus parlatorei*) inhabiting the uppermost levels of the Yungas forests. They found northern, central and southern clades that generally match those that we found, particularly for *H. riojanus*. Quiroga & Premoli [77] suggested that, as *P. parlatorei* is adapted to cool climates, the species expanded to encompass a more continuous range at lower elevations (and lower latitude) during glacial maxima, to become more fragmented at higher elevations during warmer periods. This pattern is opposite that expected for frogs, expansion during warmer and more mesic episodes, but consistent with the notion that any cycling between cool and warm periods may result in genetic divergence of populations. Hensen et al. [8] investigated the genetic structure of *Polylepsis australis*, a tree endemic to mountainous regions of northwestern Argentina. Their study encompasses our study area plus the additional mountain region of Cordoba, 482 km to the southeast. They identified two distinct lineages across the geographic range of the species separating groups found (i) in the north (similar distribution to our “northern” clades), and (ii) those in the central (similar distribution to our “southern” clades) as well as southern Cordoba populations. The north to south distribution of lineages and the genetic break between lineages found at approximately 26-27°S mirrors genetic patterns that we found. Hensen et al. [8] postulate that, converse to the expected patterns from the Northern Hemisphere, during climate warming *P. australis* populations would have migrated northwards rather than southwards.

In northwestern Argentina, three potential refugia can be identified based on species distributions [78] and on the genetic divergence patterns described for of the aforementioned co-distributed taxa. Northern (~22°S) and southern (~27°S) refugia have been proposed previously based on high species richness and endemism [77] and current highly mesic climate. Our results suggest a third refugium, situated in the Sierra de Santa Barbara (~24.5°S) given the genetically distinct lineages associated with this area and the current discontinuous distribution of Yungas forests and possible persistence of mesic habitats in the past.

## CONCLUSION

Our data support a vicariance model of diversification for both species rather than the gradient divergence model across elevations. Guarnizo et al. [6] also rejected the gradient divergence model using neutral genetic markers within the tree frog *Dendropsophus labialis* in the northern Andes of Columbia [6]. However, traits important in local adaptation may respond faster to strong selection pressures associated with environmental gradients, such as those associated with elevation, than neutral genetic markers [12]. Thus, we cannot discount a role for environmental gradients in diversification, as we may be unable to detect such a pattern during the early stages of speciation. Additional research combining data from genomics and transcriptomics, morphology, acoustics and life history may further clarify the processes of diversification in these environmentally complex landscapes.

Our study supports an important role of mountain habitats in promoting and potentially maintaining biodiversity (i) through fragmentation and reconnection among populations, providing opportunities for differentiation and possibly speciation, and (ii) because of the preservation of many different lineages that are able to shift to different elevations to avoid unfavorable habitats (and possibly extinction). We also show that the southern terminus of the Andean montane forest system has been particularly dynamic over the last few million years, producing strikingly divergent and spatially coincident evolutionary lineages within long-recognized species.

## ACKNOWLEDGEMENTS

Provincial governments and National Parks of Argentina kindly granted collection and export permits. We thank all the people who assisted with field collections, especially M. Vaira. We are grateful to many laboratory assistants, and especially to M.-A. Lachance for providing expertise and laboratory space. Thank you to J. Padial and D. Ferraro for additional samples. V. Friesen and N. Keyghobadi provided insightful comments. Funding was provided by NSERC Discovery grants to Lougheed and Handford; an NSERC scholarship, an OGSST scholarship, a Sigma Xi Grant in Aid of Research, a Gaige Award from the ASIH, and various University of Western Ontario awards to Koscinski.

---

**S1 Text. Supplementary methods on PCR and sequencing steps, and details on all phylogenetic analyses.**

**S1 Fig. Bayesian skyline plots for *H. riojanus* (top) and *P. borellii* (bottom).**

The thick solid line is the median estimate, and the pale grey lines show the 95% HPD limits. Both species show population growth less than 1 Ma.

**S1 Table. Voucher specimens of *Pleurodema borellii* collected during the study.**

Typically only two individuals per population were kept as voucher specimens, the remaining individuals were toe clipped and released. Some specimens have not been assigned voucher numbers yet and are listed as pending.

**S2 Table. Results from mismatch distributions are shown for each lineage of *Pleurodema borellii*.**

Timing of population expansion estimated by τ = 2μt, where μ is the mutation rate per generation per gene and t is time; mismatch distribution is not significantly different from the sudden expansion model when p>0.05. Refer to Fig 5 for mtDNA clade numbers.

